# Notch Signaling and Fluid Shear Stress in Regulating Osteogenic Differentiation

**DOI:** 10.1101/2022.07.30.502120

**Authors:** Yuwen Zhao, Kiarra Richardson, Rui Yang, Zoe Bousraou, Yoo Kyoung Lee, Samantha Fasciano, Shue Wang

## Abstract

Osteoporosis is a common bone and metabolic disease that is characterized by bone density loss and microstructural degeneration. Human bone marrow-derived mesenchymal stem cells (hMSCs) are multipotent progenitor cells with the potential to differentiate into various cell types, including osteoblasts, chondrocytes, and adipocytes, which have been utilized extensively in the field of bone tissue engineering and cell-based therapy. Although fluid shear stress plays an important role in bone osteogenic differentiation, the cellular and molecular mechanisms underlying this effect remain poorly understood. Here, an LNA/DNA nanobiosensor was exploited to monitor mRNA gene expression of hMSCs that were exposed to physiologically relevant fluid shear stress to examine the regulatory role of Notch signaling during osteogenic differentiation. First, the effects of fluid shear stress on cell viability, proliferation, morphology, and osteogenic differentiation were investigated and compared. Our results showed shear stress modulates hMSCs morphology and osteogenic differentiation depending on the applied shear and duration. By incorporating this LNA/DNA nanobiosensor and alkaline phosphatase (ALP) staining, we further investigated the role of Notch signaling in regulating osteogenic differentiation. Pharmacological treatment is applied to disrupt Notch signaling to investigate the mechanisms that govern shear stress induced osteogenic differentiation. Our experimental results provide convincing evidence supporting that physiologically relevant shear stress regulates osteogenic differentiation through Notch signaling. Inhibition of Notch signaling mediates the effects of shear stress on osteogenic differentiation, with reduced ALP enzyme activity and decreased Dll4 mRNA expression. In conclusion, our results will add new information concerning osteogenic differentiation of hMSCs under shear stress and the regulatory role of Notch signaling. Further studies may elucidate the mechanisms underlying the mechanosensitive role of Notch signaling in stem cell differentiation.

## Introduction

Osteoporosis is a systemic metabolic bone disease characterized by reduced bone formation in the bone marrow space, which leads to bone mass loss and microstructural degeneration [1]. In the United States, it is estimated that ∼ 10 million people have osteoporosis and more than 34 million are at risk [2; 3]. It is also estimated that osteoporosis causes more than 9 million fractures annually worldwide [2]. In recent years, the cost of treating osteoporosis is increasing due to the increased aged population and space travel, causing challenges to public health care. In space, the reason for developing osteoporosis is mainly related to low (micro- to zero-) gravity conditions, with possible contributions of cosmic ray radiation [4]. For example, bone density loss occurs in the weightless environment of space due to the lack of gravity force. Thus, the bone no longer needs to support the body against gravity. Astronauts lose about 1% - 2% of their bone mineral density every month during space travel. Osteoporosis is one of the major consequences of long-duration spaceflights in astronauts, seriously undermining their health [5]. Currently, the autologous bone graft is the “gold standard” approach to restoring large bone defects with bone loss, where a piece of bone is taken from another body site, and transplanted into the defect [6]. However, the availability of donated bone and the necessity of an invasive and expensive surgery limited its application. Another approach to treat osteoporosis is to stimulate osteogenesis or inhibit bone resorption through drug-based agents, i.e., bisphosphonates [7]. However, drug-based agents are limited due to their side effects and lack of capability of regaining the lost bone density. Thus, there is an urgent need for alternative therapeutic approaches for osteoporosis, especially therapies that are able to counteract bone mass loss, which is crucial for aged populations and astronauts that are needed for prolonged space missions.

Human bone marrow-derived mesenchymal stem cells (hMSCs) are ideal candidates for cell-based therapies for bone tissue engineering and regenerative medicine due to their multipotency. Under mechanical or chemical stimulation, hMSCs can be induced to differentiate into various lineages, including osteoblasts (bone), neuroblasts (neural tissue), adipoblasts (fat), myoblasts (muscle), and chondroblasts (cartilage) [8]. Moreover, the fate commitment and differentiation of hMSCs is closely controlled by the local mechanical and chemical environment that maintains a balance between osteogenic differentiation and adipogenic differentiation. Reduced osteogenic differentiation and increased adipogenic differentiation might lead to osteoporosis. Although the differentiation capacity of hMSCs has been demonstrated, the mechanisms that control their plasticity remain poorly understood, especially how hMSCs can be differentiated into osteoblasts and make bones. It is believed that mechanical stimulation impacts hMSCs osteogenic differentiation. Over the last few decades, unremitting efforts have been devoted to understanding biochemical signals that regulate hMSCs commitment. Based on these efforts, a number of chemical stimuli (e.g., small bioactive molecules, growth factors, and genetic regulators) have been identified in regulating hMSCs lineage commitment, including bone morphogenetic protein (BMP), Wnt, and Notch signaling [9; 10; 11]. Since the last decade, the effects of physical/mechanical cues of the microenvironment on hMSCs fate determination have been investigated extensively. For instance, several studies provide evidence that mechanical cues, including shear, stiffness and topography, and electrical stimulation, and acoustic tweezing cytometry (ATC) [12; 13], both direct and indirect, play important roles in regulating stem cell fate. Moreover, it had been shown that ECM and topography enhance hMSCs osteogenic differentiation by cellular tension and mechanotransduction of YAP activity [14; 15; 16; 17]. Although these studies have made significant progress in understanding the stimuli that regulates hMSCs differentiation, the fundamental mechanism of osteogenic differentiation remains uncharacterized. Particularly, the interaction of biophysical factors and biochemical signals is obscure. Thus, understanding the interaction of biophysical and chemical signals in osteogenic differentiation may provide new insights to improve our techniques in cell-based therapies and organ repair.

Osteogenic differentiation is a dynamic process and involves several significant signaling pathways, including YAP/TAZ, Notch, and RhoA signaling [18; 19]. It has been shown that fluid shear force, including that encountered in microgravity models, regulates *in vitro* osteogenic differentiation of MSCs [20; 21; 22; 23]. For example, it has been shown that physiologically relevant fluid-induced shear stress of 3-9 dynes/cm^2^ could be conducive to cell conditioning, and assist in promoting genes [24; 25; 26]. It is also reported that hMSCs were able to differentiate into endothelial cells and activate interstitial cells deeper when exposed to physiologically relevant steady fluid-induced shear stress (4-5 dynes/cm^2^) [27]. Although current studies revealed shear stress could enhance osteogenic differentiation, the involvement of Notch signaling in shear stress induced osteogensis is not clear.

Here, we exploited a double-stranded locked nucleic acid/DNA (LNA/DNA) nanobiosensor to elucidate the regulatory role of Notch signaling during osteogenic differentiation of hMSCs that were exposed to physiologically relevant shear stress (3-7 dynes/cm^2^). The effects of fluid shear stress on hMSCs proliferation and osteogenic differentiation were first investigated and compared under different levels of fluid shear stress. The phenotypic behaviors, including cell morphology, proliferation, and differentiation, were compared and characterized. We further detected Notch 1 ligand Delta-like 4 (Dll4) gene expression by incorporating this LNA/DNA nanobiosensor with hMSCs imaging during osteogenic differentiation. Finally, we examined the role of Notch signaling in regulating osteogenic differentiation of hMSCs that are under shear stress. Pharmacological administration is applied to disrupt Notch signaling to investigate the cellular and molecular mechanisms that govern osteogenic differentiation. Our experimental results provide convincing evidence supporting that physiologically relevant shear stress regulates osteogenic differentiation through Notch signaling. Inhibition of Notch signaling will mediate the effects of shear stress on osteogenic differentiation, with reduced alkaline phosphatase (ALP) enzyme activity and decreased Dll4 mRNA expression. In conclusion, our results will add new information concerning osteogenic differentiation of hMSCs under shear stress and the involvement of Notch signaling. Further studies may elucidate the mechanisms underlying the mechanosensitive role of Notch signaling in stem cell differentiation.

## Materials and Methods

### Cell culture and reagents

Human Bone Marrow Derived Mesenchymal Stem Cells (hMSCs) were acquired from Lonza, which were isolated from normal (non-diabetic) adult human bone marrow withdrawn from bilateral punctures of the posterior iliac crests of normal volunteers. hMSCs were cultured in mesenchymal stem cell basal medium MSCBM (Catalog #: PT-3238, Lonza) with GA-1000, L-glutamine, and mesenchymal cell growth factors (Catalog #: PT-4105, Lonza). Cells were cultured in a tissue culture dish at 37 °C and 5% CO_2_ in a humidified incubator. Cells were maintained regularly with medium change every three days and passaged using 0.25 % EDTA-Trypsin (Invitrogen). hMSCs from passage 2-7 were used in the experiments. For osteogenic induction studies, hMSCs were seeded at a concentration of 400 cells/mm^2^ with a volume of 500 μL basal medium in 24 well-plates. Once the cells reach 80% confluency, for the control group, cells were maintained in basal medium. For induction group, the basal medium was replaced with osteogenic differentiation medium (Catalog #: PT-3002, Lonza). Osteogenic differentiation medium was changed every two days. For studying Notch signaling, hMSCs were treated with 20 μM γ-secretase inhibitor DAPT (Sigma Aldrich) after osteogenic induction. It is noted that DAPT treatment was performed daily. Images were taken after 3 days and 5 days of osteogenic induction, respectively.

### Design of LNA probe

An LNA detecting probe is a 20-base pair nucleotide sequence with alternating LNA/DNA monomers that is complementary to target mRNA sequence with a 100% match. For target mRNA detection, a fluorophore (6-FAM) was labeled at the 5’ end of the LNA probe for fluorescence detection. The design process of the LNA probe for mRNA detection was reported previously.[28; 29; 30; 31; 32; 33] Briefly, the target mRNA sequence was first acquired from GeneBank. A 20-base pair nucleotide sequence was selected and optimized using mFold server and NCBI Basic Local Alignment Search Tool (BLAST) database. A quencher probe is a 10-base pair nucleotide sequence with LNA/DNA monomers that is complementary to the 5’ end of the LNA detecting probe. An Iowa Black RQ fluorophore was labeled at the 3’ end of the quencher probe. The Dll4 LNA detecting probe was designed based on target mRNA sequences (5’-3’: +AA +GG +GC +AG +TT +GG +AG +AG +GG +TT). The LNA detecting probe and quencher sequence were synthesized by Integrated DNA Technologies Inc. (IDT).

### Preparation of double-stranded LNA probe

To prepare the LNA/DNA nanobiosensor, the LNA detecting probe and quencher probe were initially prepared in 1x Tris EDTA buffer (pH=8.0) at a concentration of 100 nM. The LNA probe and quencher were mixed at the ratio of 1:2 and incubated at 95 °C in a dry water bath for 5 minutes and cooled down to room temperature over the course of 2 h. Once cooled down, the prepared LNA probe and quencher mixer can be stored in a refrigerator for up to 7 days. For mRNA detection, the prepared double-stranded LNA/DNA probe was then transfected into hMSCs using Lipofectamine 2000 following manufacturers’ instructions. mRNA gene expression can thus be evaluated by measuring the fluorescence intensity of hMSCs transfected with LNA/DNA probes.

### Simulation of orbital shear stress

hMSCs were exposed to 30/60 RPM orbital shear stress using a low-speed orbital shaker (Corning LSE, 6780-FP, orbit, 1.9cm, speed range, 3-60 rpm). The orbital shear was applied to hMSCs after osteogenic induction for 6 hours per day or continuously for a total of 3 and 5 days. The orbital shaker was placed inside the incubator to maintain cell environment. The orbital shear stress was calculated using the following equation:

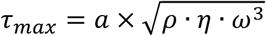

Where *τ*_*max*_ is near-maximal shear stress, *a* is the orbital radius of rotation, *ρ* is the density of cell culture medium, *η* is the dynamic viscosity of the medium, *ω* is the angular velocity and *ω* = 2*πf. f* is the frequency of rotation (revolution per second).

### Cell proliferation and viability

To evaluate the effects of applied orbital shear stress on hMSCs proliferation and viability, a cell proliferation and viability reagent (Cell Counting kit-8, cck-8 assay, Sigma Aldrich) was utilized following the manufacturers’ instructions. First, hMSCs were seeded in three flat-bottom 96-well tissue culture well plates with the density of 2000 cells/well with the volume of 100 μL basal culture medium. After 24 hr of incubation to allow cell attachment, two 96-well plates were placed on orbital shaker. Out of these two well-plates, one well plate was kept on the orbital shaker to experience continuous orbital shear stress for 3 or 5 days, the other well plates was kept on the orbital shaker for 6 hr per day for a duration of 3 or 5 days. The third 96-well plate was kept in static condition in the incubator for comparison. Cell viability was evaluated after 3 or 5 days of applying shear stress. After applying shear, CCK-8 reagents were added to each well and incubated for 4 hr. The absorbance of each sample was measured at 450 nm and compared using a fluorescence microplate reader (BioTek, Synergy 2).

### Live/Dead viability staining

The hMSCs viability after orbital shear was evaluated using live/dead viability assay (ThermoFisher). hMSCs were stained using propidium iodide (PI, 10 μg/mL), a fluorescent agent that binds to DNA by intercalating between the bases with little or no sequence preference. The cell nucleus was stained using Hoechst 33342 for 30 minutes at the concentration of 20 μM. After staining, hMSCs were washed three times with 1x PBS to remove extra dye. hMSCs were then imaged using Texas Red (535/617 nm) and DAPI (360/460 nm) filters on the ZOE image station.

### Staining

To quantify hMSCs osteogenic differentiation, alkaline phosphatase enzyme activities were evaluated and measured by using two ALP staining assays, AP live staining (ThermoFisher) and ALP staining kit (for fixed cells, Sigma-Aldrich). For fixed cells, the staining solution was first prepared by mixing Fast Red Violet solution, Naphthol AS-BI phosphate solution and water at a ratio of 2:1:1. Next, hMSCs were fixed using 4% cold-Paraformaldehyde (PFA) for 2 minutes which enable the maintenance of the ALP activation. After fixation, the PFA was aspirated without wash. The staining solution was then added to the fixed cells for 15 minutes under room temperature and protected from light. The cells were then washed three times with 1x PBS, 15 minutes each time, before taking images. For AP live staining, hMSCs were stained using AP live stain at the concentration of 10X stock solution for 30 minutes according to manufacturers’ instructions. After staining, hMSCs were washed twice using basal medium. Images were captured after 30 minutes of staining. For F-actin staining, hMSCs were first fixed with 4% PFA solution for 10 minutes before being permeabilized and blocked with the PBST solution (PBS + 0.5% Triton + 1% BSA) for 1 hr. After wash with 1x PBS three times, hMSCs were incubated with phalloidin (1:30) for 1 hr at room temperature. The cells were then washed three times using 1x PBS, before imaging.

### Imaging and Statistical Analysis

Images were captures using ZOE Fluorescent Cell Imager with an integrated digital camera (BIO-RAD) or Nikon TE 2000 with a Retiga R1 monochrome CCD Camera. For comparison, all the images were taken with the same setting, including exposure time and gain. Data collection and imaging analysis were performed using NIH ImageJ software. To quantify Dll4 mRNA and ALP enzyme activity, the mean fluorescence intensity of each cell was measured. The background noise was then subtracted. All the cells were quantified in the same field of view and at least five images for each condition were quantified. All experiments were repeated at least three times and over 100 cells were quantified for each group. Results were analyzed using independent, two-tailed Student *t*-test in Excel (Microsoft). P < 0.05 was considered statistically significant.

## Results

### Design LNA/DNA nanobiosensor for mRNA detection

To investigate the involvement of Notch signaling in osteogenic differentiation of hMSCs that were exposed to shear stress, we utilized an LNA/DNA nanobiosensor for mRNA gene expression analysis. The LNA/DNA nanobiosensor is a complex of an LNA detecting probe and a quencher, **Fig. 1A**. The LNA detecting probe is a 20-base pair single stranded oligonucleotide sequence with alternating LNA/DNA monomers, which are designed to be complementary to the target mRNA sequence. The LNA nucleotides are modified DNA nucleotides with higher thermal stability and specificity [34]. A fluorophore (6-FAM (fluorescein)) was labeled at the 5’ end of the LNA detecting probe for mRNA detection. Design, characterization, and optimization of LNA/DNA nanobiosensor have been reported previously [28; 30; 33]. Briefly, the LNA probe will bind to the quencher spontaneously to form a LNA - quencher complex. Due to their close physical proximity, the fluorophore at the 5’ end of the LNA probe is quenched by quencher due to its quenching ability [35]. After it is internalized by cells and in the presence of the target mRNA sequence in the cytoplasm, the LNA probe is thermodynamically displaced from the quencher and binds to specific target mRNA sequences, which permits the fluorophore to reacquire fluorescence signal, **Fig. 1B**. This displacement is due to the larger difference in binding free energy between LNA probe to target mRNA versus LNA probe to quencher. Thus, the fluorescence intensity of individual cells containing LNA/DNA nanobiosensor can serve as a quantitative measurement of the amount of target mRNA in each cell. In this study, hMSCs were transfected with the LNA/DNA nanobiosensor prior to osteogenic induction. The mRNA expression at the single cell level was clearly evident, **Fig. 1C**.

**Fig. 1.**
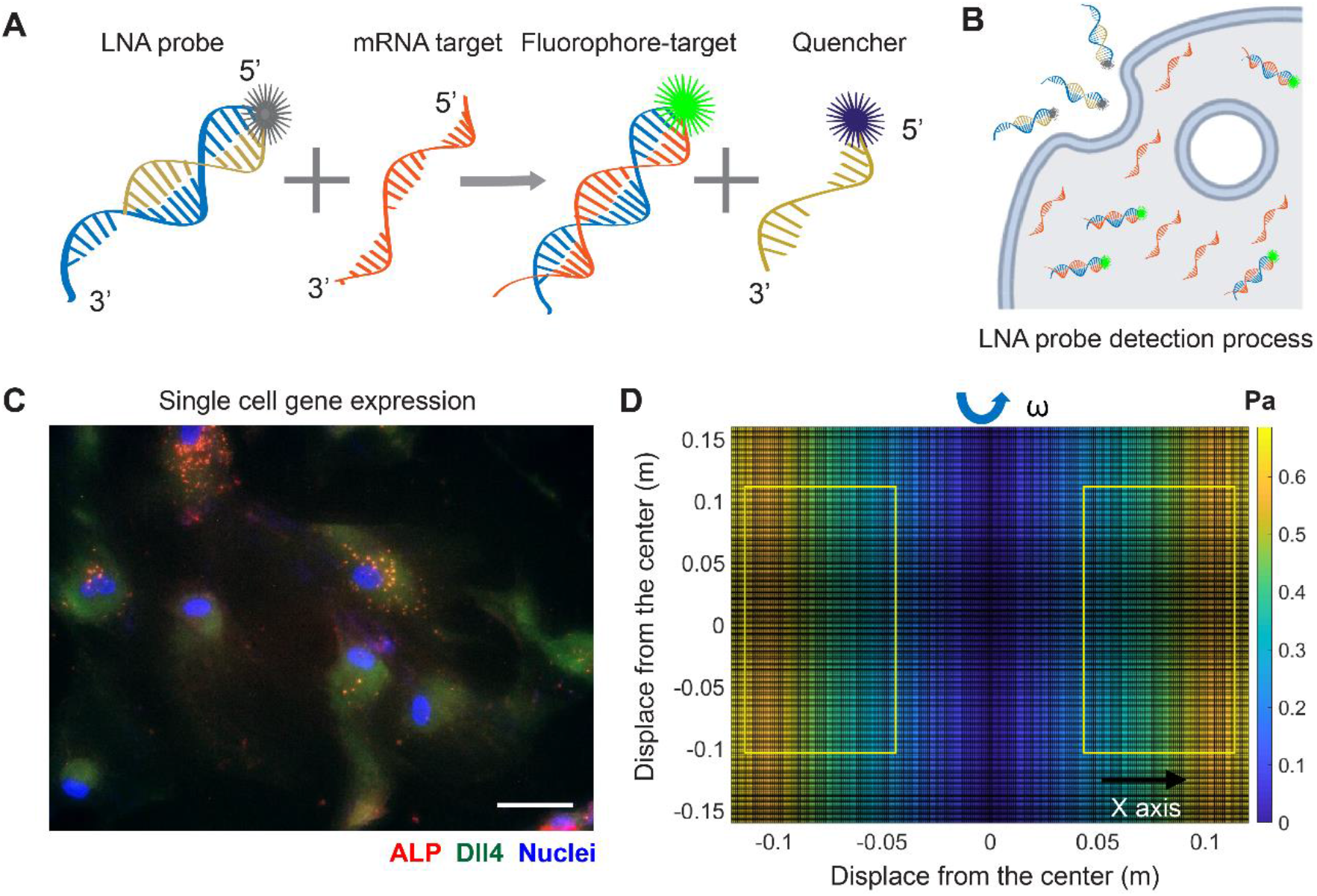
LNA/DNA nanobiosensor for single cell gene expression analysis in living cells. **(A)** Schematic illustration of LNA/DNA nanobiosensor for mRNA detection. Briefly, the LNA/DNA nanobiosensor is a complex of LNA donor and quencher probe. The fluorophore at the 5’ of LNA donor probe is quenched due to close proximity. In the presence of target mRNA sequence, the LNA donor sequence is displaced from the quencher to bind to the target sequence, allowing the fluorophore to fluorescence. **(B)** Schematic illustration of cellular endocytic uptake of LNA/DNA nanobiosensor by cells for intracellular gene detection. **(C)** Representative fluorescence image of Dll4 mRNA expression, ALP expression in hMSCs using LNA/DNA nanobiosensor. Green: Dll4 mRNA; red: ALP; blue: Nuclei. Scale bar: 100 μm. **(D)** Simulated distribution of orbital shear stress. Yellow labeled rectangles indicate the location of well-plates. The estimated shear stress were in the range of 0.3 ∼ 0.6 Pa.

### Simulation of orbital shear stress and analysis

To evaluate the effects of physiologically relevant shear stress on osteogenic differentiation, the shear stress was estimated using Strokes’ second problem, which concerns a plate oscillating along one axis in the plane of the plate, with a liquid above it. Although the orbital shaker does not produce uniform laminar shear stress on seeded cells, most of the cells were exposed to near-maximum shear that is calculated as: 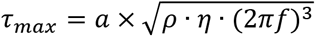, where *a* is the orbital radius of rotation. The density of hMSCs culture medium is ∼1.015 × 10^3^ kg/m^3^, the dynamic viscosity is 0.958 × 10^−4^ kg/m.s [36]. Since the cells in different wells were placed at different locations on the shaker, the shear stress is slightly different. Thus, we simulated the distribution of the shear stress over the shaker platform. Since the orbital shaker shakes along one axis (y), the shear stress along the y axis is the same. At 30 RPM, the orbital shear stress was simulated, as shown in **Figure 2A**. The maximum shear stress is approximately 0.7 Pascal (7.1 dyne/cm^2^), which is on the edge of the shaker. At the center of the shaker, the shear stress is zero. The well-plates with the dimensions of 120 mm x 85 mm were placed on the shaker, labeled in **Fig. 1D**. Thus, the applied shear stress to different wells ranges from 3 dyne/cm^2^ to 7 dyne/cm^2^, which are similar to the values reported by others [37; 38; 39].

**Fig. 2.**
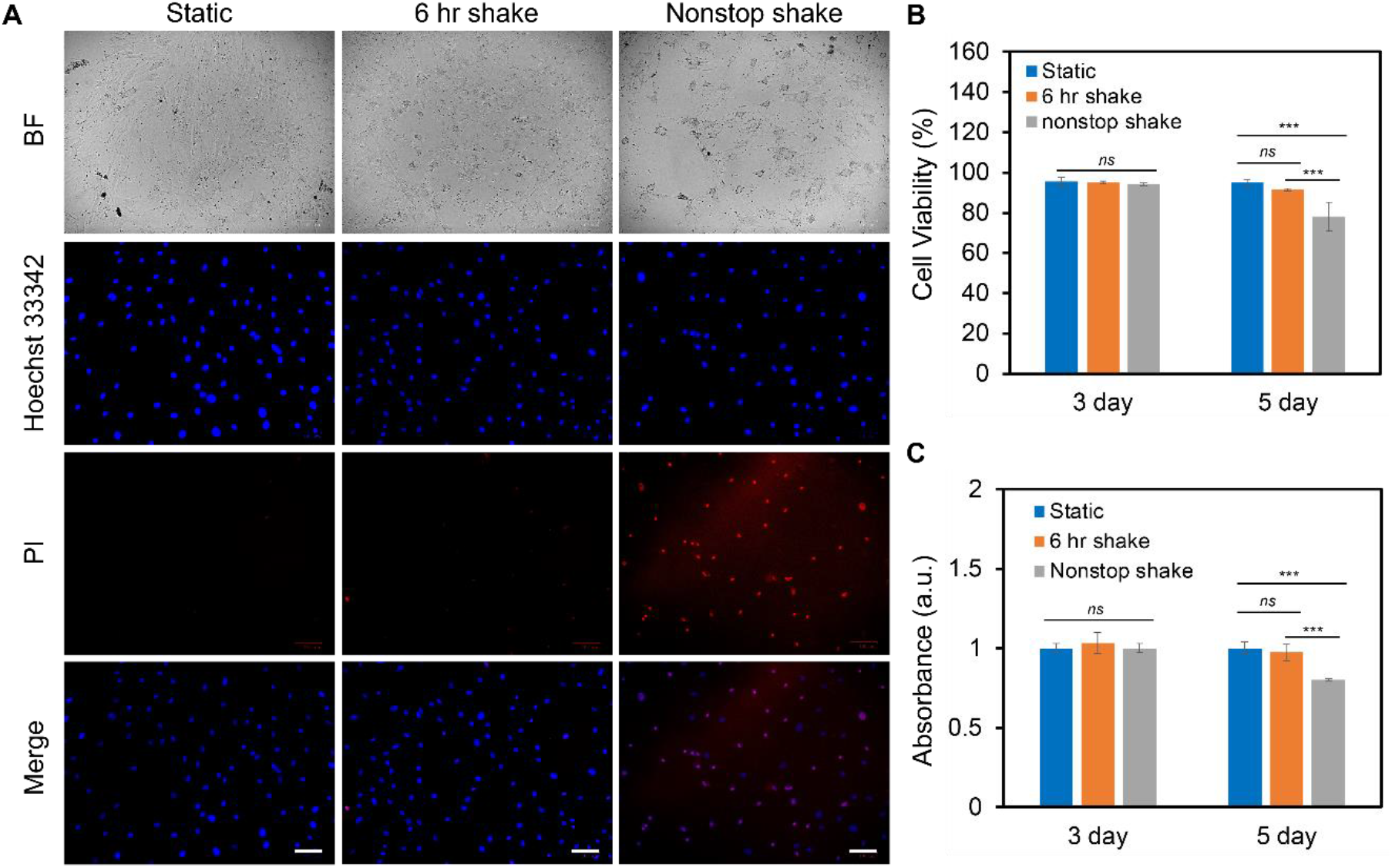
Effects of shear stress on cell viability and proliferation. **(A)** Representative bright field and fluorescence images of hMSCs after 5 days of culture with the speed of 20 RPM under static, 6 hr shake, and nonstop shake conditions, respectively. Static: cells were placed in a CO_2_ incubator without shear; 6 hr shake: cells were placed on the orbital shaker for 6 hrs per day; nonstop shake: cells were placed on the orbital shaker without stop. Samples were stained with propidium iodide (PI, red), and Hoest 33342 (blue), respectively. Scale bar: 100 μm. **(B)** Comparison of cell viability of hMSCs after 3 days and 5 days of culture under three different conditions, respectively. Cell viability was calculated as: # of live cells per field/total # of cells per field × 100%. Data represents over 500 cells in each group and expressed as mean± s.e.m. (n=4, *ns*, not significant, ***, P<0.001, **, P<0.01). **(C)** Comparison of the proliferation of hMSCs cultured in different conditions. Data were acquired using a cck-8 assay and the absorbance at 450 nm was compared. Data are expressed as mean± s.e.m. (n=4, *ns*, not significant, ***, P<0.001, **, P<0.01)

### Fluid Shear Stress modulates hMSCs proliferation and viability

In order to study the effects of different levels of shear stress on cell proliferation and viability, three groups of experiments were designed and compared: static condition, 6 hr shake, and nonstop shake. For static condition, cells were placed in the humidified CO_2_ incubator without applying shear; for the 6 hr shake, cells were applied shear stress for 6 hr per day for a total duration of 3 and 5 days; for the nonstop shake group, cells were applied orbital shear stress without a stop for a total of 3 and 5 days. Two different levels of shear stress were investigated: low fluid shear stress and high fluid shear stress. The low fluid shear stress were defined as the shear stress that is physiologically relevant with a range of 1-9 dynes/cm^2^; while high fluid shear stress is double the magnitude of low fluid shear stress (9 - 20 dynes/cm^2^). The cell viability and proliferation were evaluated using live/dead cell assay and cell counting kit (cck-8) assay after 3 days and 5 days, respectively. Under low fluid shear stress, the cell viability and proliferation were evaluated and compared, **Fig. 2. Fig. 2A** shows the bright field and fluorescent images of hMSCs after 5 days of shear stress under different groups. It is evident that the number of dead cells increased when hMSCs were exposed to continuous shear for 5 days. We further quantified the effects of shear stress on cell viability and proliferation. The cell viability was calculated as: # of live cells per field / # of total cells per field × 100%. After applying shear for 3 days, the cell viability and proliferation of hMSCs under shear stress did not show a significant difference compared to hMSCs in the static condition, left panel of **Fig. 2B-2C**. After 5 days, hMSCs under continuous shear stress showed significantly reduced cell viability and proliferation, with a 21.5% decrease in cell viability and a 19.8% decrease in proliferation compared to the cells in the static condition, right panel of **Fig. 2B-2C**. It is noted that after applying shear stress for 5 days with 6 hrs per day, the cell viability and proliferation of hMSCs did not show a significant difference compared to the hMSCs that were in the static condition. Furthermore, we studied the effects of high fluid shear stress (9 - 20 dynes/cm^2^) on hMSCs viability and proliferation, **Fig. S1**. For the hMSCs that were exposed to high shear stress for 3 days, the cell viability was decreased by 55% for the nonstop shake group. After 5 days of applying shear stress, the number of dead cells in both the 6 hr shake and nonstop shake groups increased significantly, **Fig. S1A**. Moreover, compared to hMSCs in the static condition, the cell viability was decreased by 14.8% and 19.2%, respectively, **Fig. S1B**. The effect of high fluid shear stress on cell proliferation has similar effects, **Fig. S1C**. After 5 days of applying shear stress, the absorbance of hMSCs under 6 hr shear and continuous shear were significantly decreased by 23.8% and 28.3%, respectively. These results revealed that shear stress modulate cell proliferation and viability is time- and speed-dependent. With high fluid shear stress, the cell viability and proliferation were decreased. With low fluid shear stress, the viability and proliferation was not affected when cells were exposed to periodic shear (6hr shear/day) instead of continuous shear (nonstop shear). In summary, for hMSCs under low fluid shear stress with 6 hr per day for 5 days, there is no significant difference in cell viability and proliferation compared to the static condition. Thus, we chose this condition (3-7 dynes/cm^2^) to avoid the effects of shear stress on cell viability and proliferation for the rest of our studies.

### Low fluid shear stress modulates hMSCs morphology

To investigate the impacts of low fluid shear stress on hMSCs morphology, we quantified and compared cell phenotypic behaviors, including cell area, cell length, cell aspect ratio, and cell perimeter with and without shear stress for 3 days and 5 days, respectively. Cells subjected to shear stress (6 hr per day) were compared to cells that were simply plated into tissue culture plates without shear (Control group). The control group provides a benchmark to account for any effects of exposing the cells to shear stress. For dynamic culture, hMSCs were exposed to shear stress (∼3-7 dyne/cm^2^) for 3 days or 5 days with 6 hours per day. After 3 days or 5 days of static or dynamic incubation, hMSCs were fixed, stained, and analyzed. **Fig. 3A-B** showed the representative bright field and fluorescence images of hMSCs under static conditions (**Fig. 3A**) and hMSCs that were exposed to shear stress (**Fig. 3B**), respectively. We further quantified and compared the cell area, aspect ratio, cell perimeter, and cell length, **Fig. 3C-3D** and **Fig. S2A-2B**. After 3 days of culture, the cell area, aspect ratio, perimeter, and cell length of hMSCs cultured under shear stress showed a slight increase (a 16.3% increase in cell area, a 14.9% increase in cell aspect ratio, a 18% increase in perimeter, and a 12% increase in cell length) compared to hMSCs cultured in the static condition. However, hMSCs exposed to low fluid shear stress for 5 days showed a 55% increase in cell area, a 72% increase in cell length, a 16% increase in cell aspect ratio, and a 30% increase in cell perimeter, respectively, compared to hMSCs cultured under static conditions, **Fig. 3C-3D** and **Fig S2A-2B**. These results indicate that hMSCs are sensitive to low fluid shear stress with significant morphology changes. This finding is consistent with previously reported studies [37; 40].

**Fig. 3.**
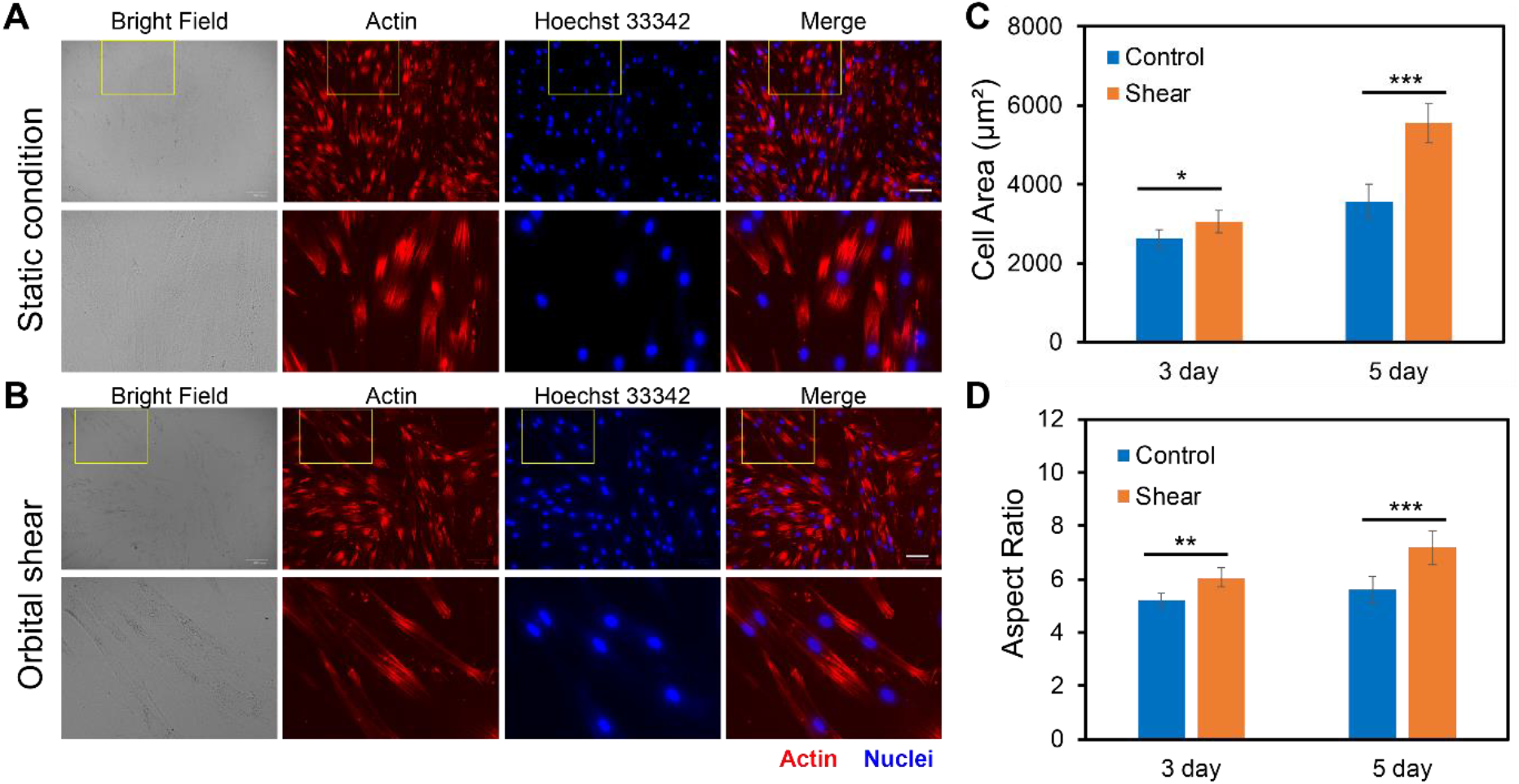
Effects of orbital shear stress on hMSCs morphology change. Representative bright field and fluorescence images of hMSCs under static condition **(A)** and exposed to shear stress **(B)**. The bottom panel showed the enlarged area of a yellow rectangle in the upper panel. hMSCs were exposed to orbital shear for 6 hours per day for 5 days. Samples were stained with F-actin (red; by phalloidin), and nuclei (blue; by Hoest 33342), respectively. Scale bar: 100 μm. Quantification of observed cell area **(C)** and cell aspect ratio **(D)** of hMSCs after 3 days and 5 days of exposure to orbital shear with 6 hours per day. Data represent over 100 cells in each group and are expressed as mean± s.e.m. (n=5, ***, P<0.001, **, P<0.01, *, P<0.05).

### Low fluid shear stress promotes osteogenic differentiation

We further elucidated the effects of low fluid shear stress on osteogenic differentiation by applying shear with the estimated shear stress of 3-7 dyne/cm^2^. Briefly, hMSCs were initially seeded in two well plates and cultured in the basal medium under static condition. Once the cells reached 70-80% confluency, osteogenic induction was performed and one well plate was placed on top of the orbital shaker, while the other plate was placed in the static condition without exposure to shear. After 5 days of osteogenic induction and shaking, osteogenic differentiation was evaluated and compared by measuring ALP enzyme activity, a reliable biochemical marker for early osteogenic differentiation [41]. The ALP enzyme activities of hMSCs were imaged, quantified, and compared after 5 days of osteogenic induction for both groups. F-actin and nucleus were also stained for better identification of each cell. **Fig. 4A-4B** showed representative bright field and fluorescence images of hMSCs under static condition and shear stress, respectively. The results showed that without osteogenic induction, there is a minimum green fluorescence signal, which indicates minimum ALP enzyme activity. With osteogenic induction, ALP enzyme activity was significantly increased in hMSCs under static condition and shear stress. We further quantified and compared ALP activity by measuring the mean green fluorescence intensity of ALP stained hMSCs. The fluorescence intensity was normalized for better comparison. Under the static condition, the ALP activity of hMSCs cultured in osteogenic induction medium increased by 1.8 folds compared to hMSCs cultured in basal medium. Under low fluid shear stress, the ALP activity was increased by 2.1 folds. Moreover, compared to the static condition, hMSCs exposed to shear stress showed a 15% increase ((ALP intensity of hMSCs with shear – ALP intensity of hMSCs without shear)/ALP intensity of hMSCs without shear) of ALP activity after osteogenic induction, **Fig. 4C**. We further quantified the differentiation percentage of hMSCs with and without fluid shear stress, which was calculated by the number of ALP labeled cells per field/ total number of cells per field. With low fluid shear stress, the hMSCs differentiation percentage increased to 45.51%, compared to 38.02% for hMSCs under the static condition. These results indicate that low fluid shear stress significantly enhanced osteogenic differentiation with increased ALP enzyme activity and osteogenic differentiation rate.

**Fig. 4.**
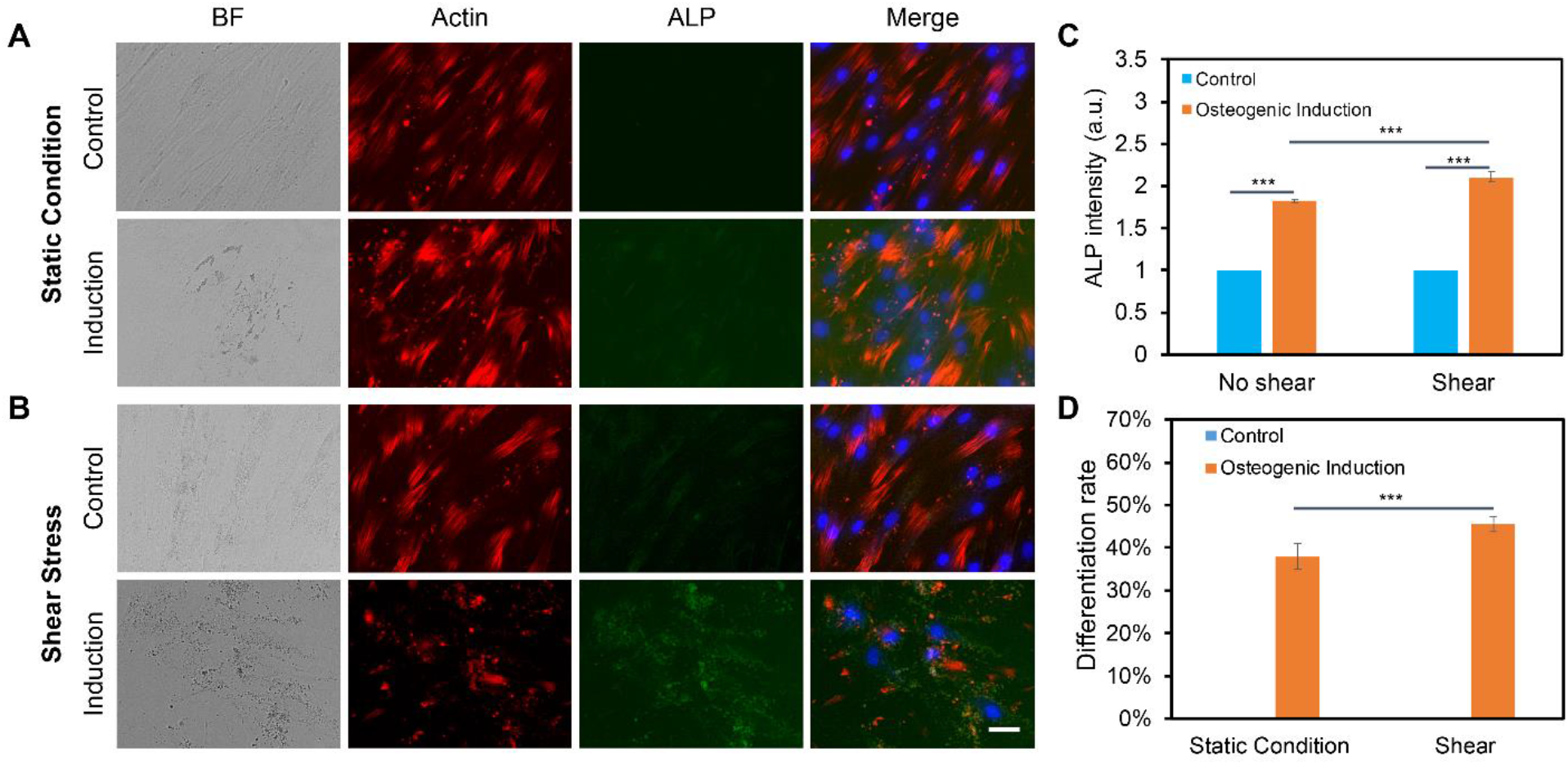
Orbital shear stress enhanced hMSCs osteogenic differentiation. Representative bright field and fluorescence images of hMSCs cultured with basal culture medium and osteogenic induction medium for 5 days under static condition **(A)**, and exposed to shear stress **(B)**, respectively. hMSCs that were exposed to orbital shear stress for 5 days with 6 hours per day. Samples were stained with ALP (green; by ALP live stain), F-actin (red; by phalloidin), and nuclei (blue; by Hoest 33342), respectively. Scale bar: 100 μm. **(C)** Fluorescence intensity of ALP activity of hMSCs with and without shear stress after 5 days of osteogenic induction compared to control group. **(D)** Osteogenic differentiation percentage with and without shear stress. Data represent over 100 cells in each group and are expressed as mean± s.e.m. (n=6, ***, P<0.001, **, P<0.01)

### Notch signaling is involved in shear stress induced osteogenic differentiation

The previous study has shown that Notch signaling is involved during hMSCs osteogenic differentiation, disruption of Notch signaling mediated ALP activity, and osteogenic differentiation efficiency [42]. Our group also recently showed that Dll4 mRNA is a molecular biomarker of osteogenic differentiated hMSCs [28]. Inhibition of Notch signaling reduces osteogenic differentiation with decreased ALP enzyme activity. However, it is obscure whether low fluid shear stress regulates osteogenic differentiation of hMSCs through Notch signaling. To better understand the involvement of Notch signaling during osteogenic differentiation, we utilized a pharmacological drug, DAPT, to perturb Notch signaling. DAPT is a γ-secretase inhibitor that blocks Notch endoproteolysis and thus serves as a Notch signaling inhibitor [43]. hMSCs were treated with DAPT at a concentration of 20 μM during osteogenic differentiation with or without shear stress to observe potential related effects. A control group was designed without osteogenic induction. The osteogenic differentiation under different treatments was evaluated and compared by measuring the mean red fluorescence intensity to examine osteogenic differentiation efficiency. **Fig. 5A-5B** and **Fig. S3A-S3B** show representative images of hMSCs under static condition and shear stress that were cultures in basal medium, induction medium, and induction medium with the treatment of DAPT, respectively. These results indicate that inhibition of Notch signaling using γ-secretase inhibitor DAPT mediated osteogenic differentiation in both static condition and shear stress. Particularly, under static condition, with the treatment of DAPT, ALP enzyme activity after 5 days of osteogenic induction was decreased by 28.8% (Fluorescent intensity with induction - Fluorescent intensity with DAPT)/ Fluorescent intensity with induction). Meanwhile, for the hMSCs exposed to low fluid shear stress, ALP enzyme activity after 5 days of induction was decreased by 18.2% with the treatment of DAPT. Interestingly, DAPT treatment for the hMSCs under shear stress has fewer effects on osteogenic differentiation, indicating low fluid shear stress rescued the inhibition effects of Notch signaling due to pharmaceutical treatment.

**Fig. 5.**
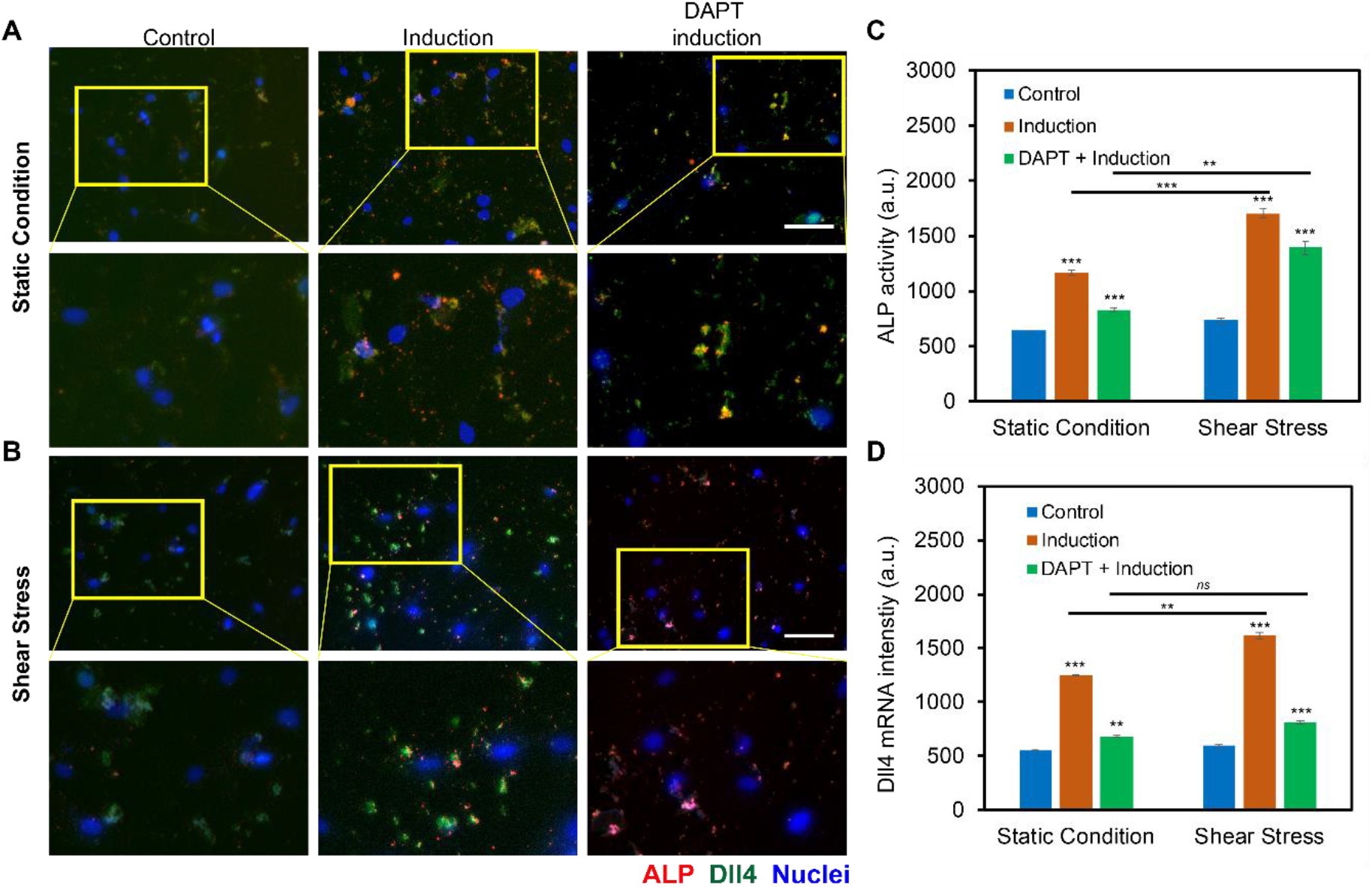
Notch signaling in regulating osteogenic differentiation of hMSCs with and without orbital shear stress. **(A)** Representative images of hMSCs in control, induction, and DAPT treatment groups without **(A)** and with **(B)** shear stress. Control: cells were cultured in the basal medium; induction: cells were cultured in osteogenic induction medium after cell seeding; DAPT: cells were treated with DAPT (20 μM) daily after osteogenic induction. Images were taken after 5 days of induction. The bottom images are enlarged areas of hMSCs in the labeled yellow rectangle. Green: Dll4 mRNA expression; red: ALP activity; blue: nucleus. Scale bar: 100 μm. **(C)** Comparison of ALP activity of hMSCs with and without shear stress under different conditions. **(D)** Mean fluorescent intensity of Dll4 mRNA expression of hMSCs after 5 days of osteogenic induction under different conditions as indicated. Error bars, s.e.m, with n = 100 - 150 cells. p-Values were calculated using a two-sample t-test with respect to control. *ns*, not significant, *, p < 0.05; **, p < 0.01; ***, p < 0.005.

To further investigate the mechanisms of Notch signaling during osteogenic differentiation that were exposed to low fluid shear stress, we examined Notch 1 ligand, Dll4 mRNA expression under static condition and shear stress with basal culture medium, induction medium, and induction medium with DAPT treatment using an LNA/DNA nanobiosensor. **Fig. 5A-5B** and **Fig. S4A-S4B** showed representative images of hMSCs under static condition and shear stress with different treatments. Dll4 mRNA expression were quantified and compared by measuring the mean green fluorescent intensity. Under the static condition, hMSCs cultured with osteogenic induction medium show a significant increase in the expression of Dll4 mRNA (∼2.26 folds increase). Meanwhile, with the treatment of γ-secretase inhibitor DAPT, a significant decrease of Dll4 mRNA (∼45.6 %) was observed compared to the osteogenic induction group, **Fig. 5D**. When exposed to low fluid shear stress, Dll4 mRNA expression of hMSCs under osteogenic induction group was increased 2.72 folds compared to hMSCs that were cultured in basal medium. The treatment of DAPT inhibited osteogenic differentiation by ∼ 50%, **Fig. 5D**. Compared to static condition, Dll4 mRNA expression was increased by 22.8% when hMSCs were cultured in osteogenic induction medium. DAPT treatment mediated the effects of shear stress on osteogenic differentiation, with only ∼16% increase of Dll4 mRNA expression. These results provide evidence that Notch signaling is involved and regulates osteogenic differentiation of hMSCs under low fluid shear stress. Low fluid shear stress upregulates Dll4 mRNA expression of hMSCs that were under osteogenic induction, indicating the involvement of Notch signaling in mechanoregulated osteogenic differentiation. Inhibition of Notch signaling mediated the effects of shear stress induced osteogenic differentiation, with reduced ALP enzyme activity and decreased Dll4 mRNA expression.

## Discussions

In this study, we investigated the role of Notch signaling in regulating osteogenic differentiation of hMSCs induced by physiologically relevant shear stress using an LNA/DNA nanobiosensor. Unlike traditional techniques for mRNA detection, this LNA/DNA nanobiosensor is capable of detecting mRNA gene expression in live cells at the single cell level without lysis or fixation. This ability enables us to track the Dll4 mRNA gene expression dynamics during osteogenic differentiation.

Notch signaling is an evolutionary well-conserved pathway that regulates cell proliferation, cell fate determination, and stem cell differentiation in both embryonic and adult organs [44; 45; 46; 47]. There are four Notch receptors (Notch1-4) and five different Notch ligands (Dll1, Dll3, Dll4, Jag1, and Jag2). In recent years, the role of Notch signaling in osteogenic differentiation has attracted researchers’ interest. Several studies showed that Notch signaling is active during osteogenic differentiation [48; 49]. Notch signaling has also been reported to control tip cell formation during angiogenesis and leader cell formation during collective cell migration [43; 50]. Recently, Xu C *et. al*. reported that Notch ligand, Dll4, could induce bone formation in male mice without causing adverse effects in other organs [51]. Notch signaling also plays an important role in controlling osteoblast and osteoclast differentiation and function, and regulates skeletal homeostasis [52]. Cao *et al*. reported that Notch receptor Notch1 and Notch ligand Dll1 are involved in osteogenic differentiation [53]. They observed that Notch1 inhibition reduced ALP activity during BMP-induced osteogenic differentiation of hMSCs *in vitro*. In contrast, it has been reported that inhibition of Notch signaling promotes adipogenic differentiation of MSCs [54], indicating that Notch involvement is lineage-dependent during MSCs differentiation. Although numerous studies have demonstrated the involvement of Notch signaling during osteogenic differentiation, it is unclear whether Notch signaling regulates osteogenic differentiation of hMSCs when exposed to physiologically relevant fluid shear stress. Here, we demonstrated that Notch signaling regulates osteogenic differentiation of hMSCs that were exposed to low fluid shear stress. We first examined the effects of shear stress on cell viability and proliferation. Our results showed shear stress regulates hMSCc viability and proliferation is time- and speed-dependent. There were minimum effects when hMSCs were exposed to low fluid shear stress (3-7 dye/cm^2^) for 6 hr per day with a duration of 5 days. We next studied the effects of shear stress on hMSCs morphology and osteogenic differentiation. The results indicate that low fluid shear stress modulates hMSCs morphology and enhances osteogenic differentiation with increased ALP enzyme activity. To elucidate the mechanisms of Notch signaling during osteogenic differentiation, we investigated Dll4 mRNA expression after 5 days of induction. Without shear stress, disruption of Notch signaling using γ-secretase inhibitor DAPT reduced ALP activity and decreased Dll4 mRNA expression. When exposed to shear stress, the effects of Notch inhibition on osteogenic differentiation were partially recovered with enhanced ALP activity and increased Dll4 mRNA expression. Overall, our results suggested that Notch signaling is involved in osteogenic differentiation and Dll4 mRNA expression was increased when hMSCs were exposed to shear stress, indicating the mechanosensitive role of Notch signaling. It is also noted that Notch signaling has been reported mechanosensitive and can be activated through laser tweezer and intercellular tension [55]. Further mechanistic studies, using 2D and 3D models, are required to elucidate the molecular and cellular processes that regulate osteogenic differentiation.

## Conclusions

In this study, an LNA/DNA nanobiosensor was exploited to detect the Dll4 mRNA gene expression profile during osteogenic differentiation of hMSCs that were exposed to physiologically relevant low fluid shear stress. We first investigated the effects of shear stress on hMSCs phenotypic behaviors including cell morphology, cell proliferation, and viability. Our results showed that high fluid shear will result in decreased cell viability and proliferation, while low fluid shear stress has minimal impacts on cell viability and proliferation. Next, we utilized an LNA/DNA nanobiosensor to monitor Dll4 mRNA expression of hMSCs during osteogenic differentiation, which enables us to identify the regulatory role of Notch signaling. Our results showed that Notch signaling regulates hMSCs osteogenic differentiation. Inhibition of Notch signaling mediates osteogenic differentiation with reduced ALP enzyme activity and Dll4 expression. We further revealed that Notch signaling is involved in shear stress induced osteogenic differentiation. During osteogenic differentiation, Dll4 mRNA expression was increased when hMSCs were exposed to low fluid shear stress, indicating the involvement of Notch signaling in mechanoregulated osteogenic differentiation. Inhibition of Notch signaling mediated the effects of shear stress induced osteogenic differentiation, with reduced ALP enzyme activity and decreased Dll4 mRNA expression. In conclusion, our results provide convincing evidence that Notch signaling regulates shear stress induced osteogenic differentiation, indicating the mechanosensitive role of Notch signaling in osteogenic differentiation. Further studies may elucidate the mechanisms underlying the mechanosensitive role of Notch signaling in regulating stem cell differentiation.

## Acknowledgment

This work is supported by NSF CAREER (CMMI: 2143151) and NASA CT Space Grant Consortium Faculty Research Grant (Award Number: P-1558). Y Zhao is supported by the Provost Graduate Fellowship. Kiarra Richardson is supported by NASA CT Student Research Grant.

## Author contributions

Y.Z and S.W conceived the initial idea of the study. Y.Z, R.Y, Z.B, K.R, Y.L, and S.F performed the experiments. Y.Z and S.W contributed to the experimental design and data analysis. Y.Z and S.W wrote the manuscript with feedback from all authors.

## Data Availability Statement

The original contributions presented in the study are included in the article/Supplementary Material, further inquiries can be directed to the corresponding authors.

