## Supplemental Materials for "Notch Signaling and Fluid Shear Stress in Regulating Osteogenic Differentiation"

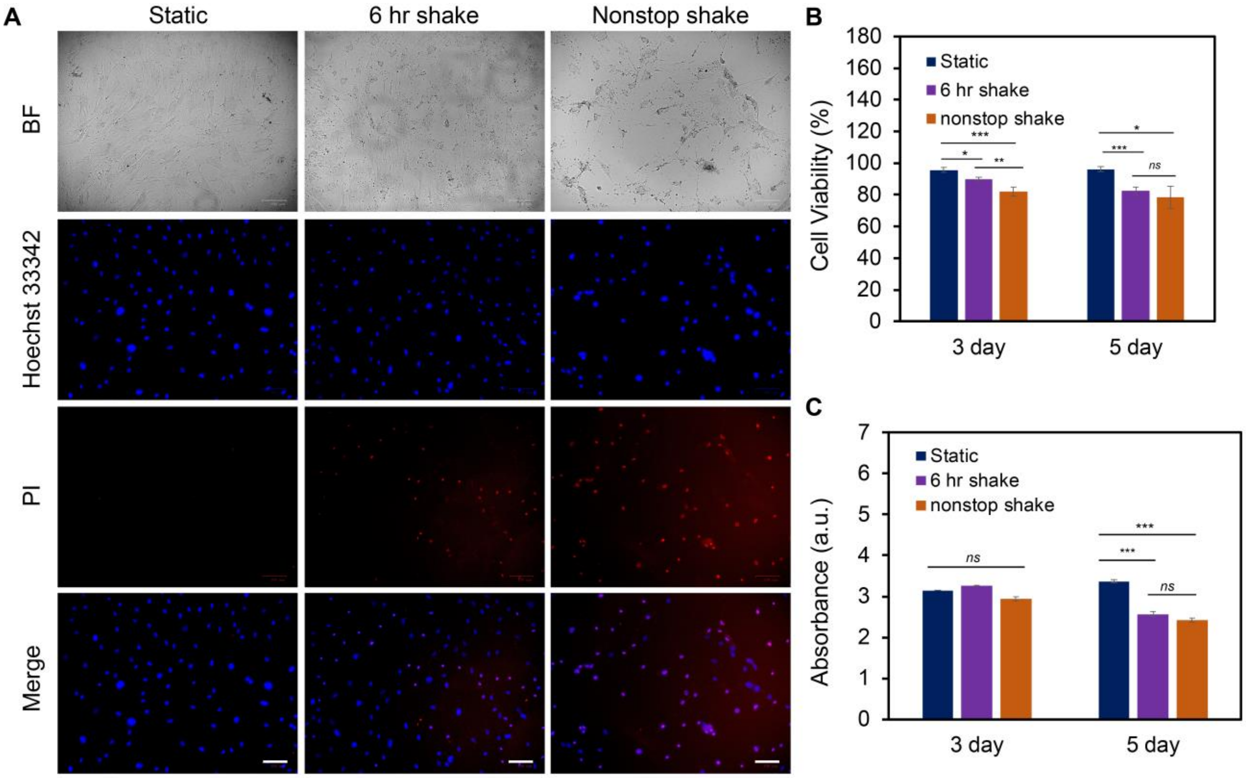


**Fig. S1**. Effects of shear stress on cell viability and proliferation. **(A)** Representative bright field and fluorescence images of hMSCs after 5 days of culture with the speed of 40 RPM under static, 6 hr shake, and nonstop shake conditions, respectively. Static: cells were placed in $CO_2$ incubator without shear; 6 hr shake: cells were placed on the orbital shaker for 6 hrs per day; nonstop shake: cells were placed on orbital shaker without stop. Samples were stained with propidium iodide (PI, red), and Hoest 33342 (blue), respectively. Scale bar: 100 µm. **(B)** Comparison of cell viability of hMSCs after 3 days and 5 days of culture under three different conditions, respectively. Cell viability was calculated as: # of live cells per field/total # of cells per field x 100%. Data represents over 500 cells in each group and is expressed as mean± s.e.m. (n=4, *ns*, not significant, ***, P<0.001, **, P<0.01). **(C)** Comparison of proliferation of hMSCs cultured in different conditions. Data were acquired using cck-8 assay and the absorbance at 450 nm was compared. Data are expressed as mean± s.e.m. (n=4, *ns*, not significant, ***, P<0.001, **, P<0.01)

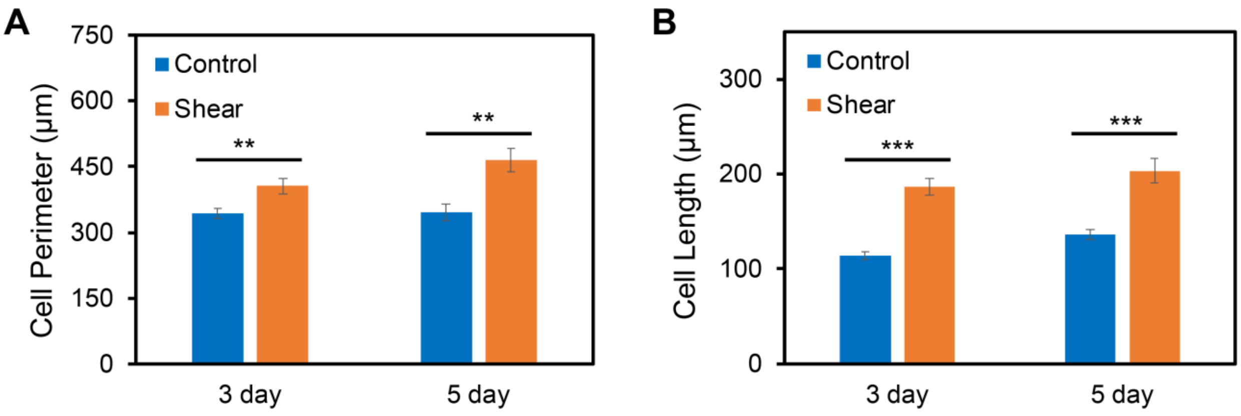


**Fig. S2**. Effects of orbital shear stress on hMSCs morphology change after 3 days and 5 days incubator, respectively. Control: cells were cultured under static conditions; shear: cells were under shear stress for 6 hrs per day for 3 or 5 days. Quantification of observed cell perimeter **(A)** and cell length **(B)** of hMSCs after 3 days and 5 days of exposure to orbital shear with 6 hours per day. Data represent over 100 cells in each group and are expressed as mean± s.e.m. (n=5, ***, P<0.001, **, P<0.01, *, P<0.05).

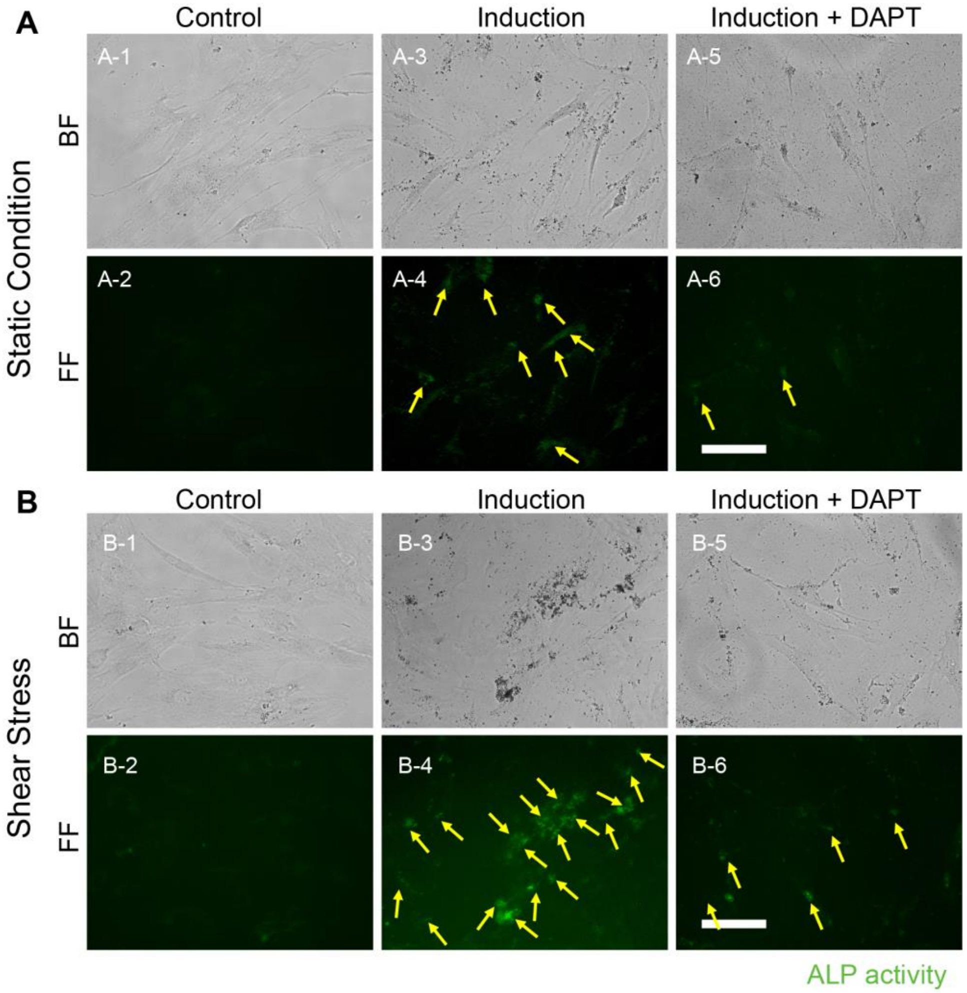


**Fig. 3**. Orbital shear stress enhanced hMSCs osteogenic differentiation. Representative bright field and fluorescence images of hMSCs cultured with basal culture medium and osteogenic induction medium for 5 days under static condition **(A)**, and exposed to shear stress **(B)**, respectively. hMSCs that were exposed to orbital shear stress for 5 days with 6 hours per day. Samples were stained with ALP (green; by ALP live stain). Scale bar: 100 μm.

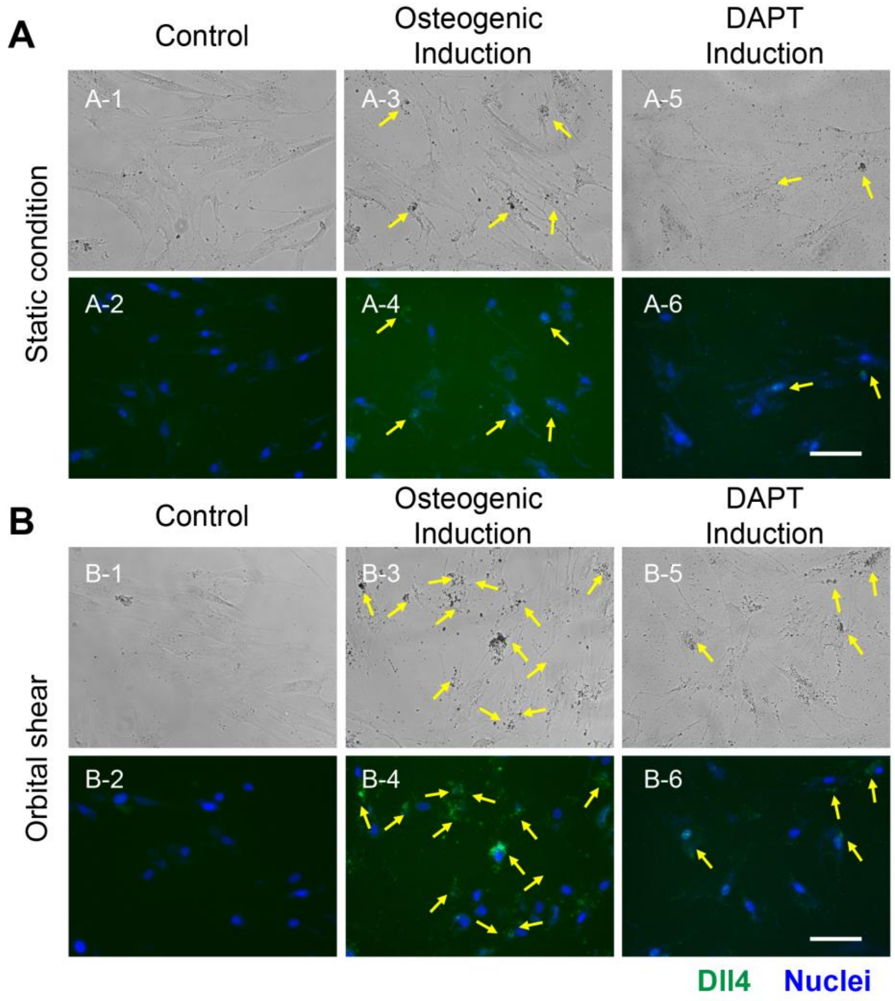


**Fig. S4**. Notch signaling in the regulation of hMSCs osteogenic differentiation. Representative bright field and fluorescence images of hMSCs with **(A)** and without **(B)** shear stress in control, induction, and DAPT treatment groups. Green: Dll4 mRNA. Blue: nucleus, respectively. Control: cells were cultured in basal medium; induction: cells were cultured in osteogenic induction medium after cell seeding; DAPT: cells were treated with DAPT (20 μM) daily after osteogenic induction. Images were taken after 5 days of induction. Scale bar: 100 μm.

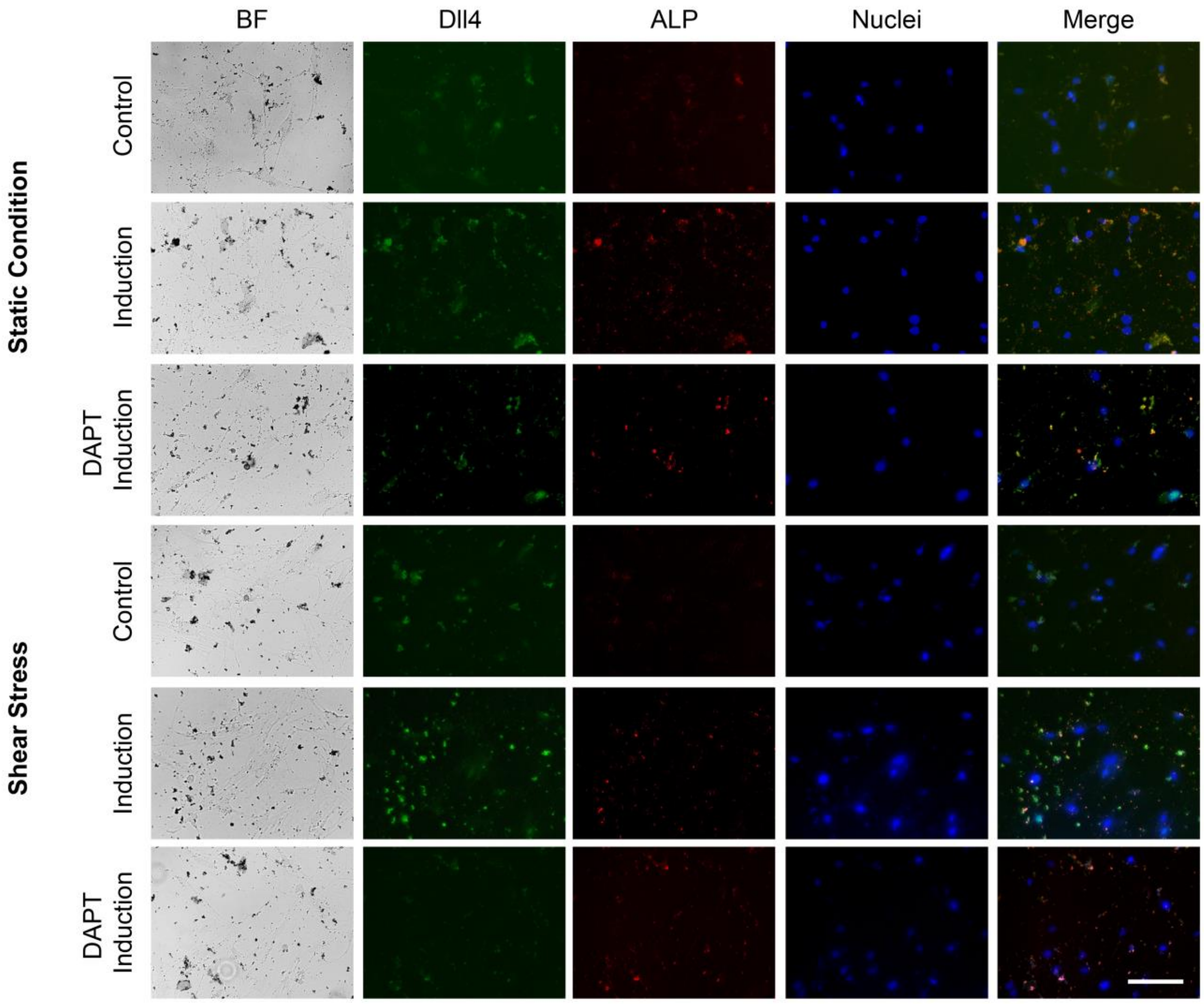


**Fig. S5**. Notch signaling in regulating osteogenic differentiation of hMSCs with and without orbital shear stress. Representative images of hMSCs in control, induction, and DAPT treatment groups without and with shear stress at separate channels. Control: cells were cultured in basal medium; induction: cells were cultured in osteogenic induction medium after cell seeding; DAPT: cells were treated with DAPT (20 μM) daily after osteogenic induction. Images were taken after 5 days of induction. Green: Dll4 mRNA expression; red: ALP activity; blue: nucleus. Scale bar: 100 μm.
